# Deep sequencing of B cell receptor repertoires from COVID-19 patients reveals strong convergent immune signatures

**DOI:** 10.1101/2020.05.20.106294

**Authors:** Jacob D. Galson, Sebastian Schaetzle, Rachael J. M. Bashford-Rogers, Matthew I. J. Raybould, Aleksandr Kovaltsuk, Gavin J. Kilpatrick, Ralph Minter, Donna K. Finch, Jorge Dias, Louisa James, Gavin Thomas, Wing-Yiu Jason Lee, Jason Betley, Olivia Cavlan, Alex Leech, Charlotte M. Deane, Joan Seoane, Carlos Caldas, Dan Pennington, Paul Pfeffer, Jane Osbourn

## Abstract

Deep sequencing of B cell receptor (BCR) heavy chains from a cohort of 19 COVID-19 patients from the UK reveals a stereotypical naive immune response to SARS-CoV-2 which is consistent across patients and may be a positive indicator of disease outcome. Clonal expansion of the B cell memory response is also observed and may be the result of memory bystander effects. There was a strong convergent sequence signature across patients, and we identified 777 clonotypes convergent between at least four of the COVID-19 patients, but not present in healthy controls. A subset of the convergent clonotypes were homologous to known SARS and SARS-CoV-2 spike protein neutralising antibodies. Convergence was also demonstrated across wide geographies by comparison of data sets between patients from UK, USA and China, further validating the disease association and consistency of the stereotypical immune response even at the sequence level. These convergent clonotypes provide a resource to identify potential therapeutic and prophylactic antibodies and demonstrate the potential of BCR profiling as a tool to help understand and predict positive patient responses.

## Introduction

Since the report of the first patients in December 2019 ^1,2,^ the unprecedented global scale of the COVID-19 pandemic has become apparent. The infectious agent, the SARS-CoV-2 betacoronavirus ^3,^ causes mild symptoms in most cases but can cause severe respiratory diseases such as acute respiratory distress syndrome in some individuals. Risk factors for severe disease include age, male gender and underlying co-morbidities ^4^.

Understanding the immune response to SARS-CoV-2 infection is critical to support the development of therapies. Recombinant monoclonal antibodies derived from analysis of B cell receptor (BCR) repertoires in infected patients or the immunisation of animals have been shown to be effective against several infectious diseases including Ebola virus ^5,^ rabies ^6^ and respiratory syncytial virus disease ^7^. Such therapeutic antibodies have the potential to protect susceptible populations as well as to treat severe established infections.

While many vaccine approaches are underway in response to the SARS-CoV-2 outbreak, many of these compositions include as immunogens either whole, attenuated virus or whole spike (S) protein - a viral membrane glycoprotein which mediates cell uptake by binding to host angiotensin-converting enzyme 2 (ACE2). The antibody response to such vaccines will be polyclonal in nature and will likely include both neutralising and non-neutralising antibodies. It is hoped that the neutralising component will be sufficient to provide long-term SARS-CoV-2 immunity following vaccination, although other potential confounders may exist, such as raising antibodies which mediate antibody-dependent enhancement (ADE) of viral entry ^8–10^. While ADE is not proven for SARS-CoV-2, prior studies of SARS-CoV-1 in non-human primates showed that, while some S protein antibodies from human SARS-CoV-1 patients were protective, others enhanced the infection via ADE ^11^. An alternative could be to support passive immunity to SARS-CoV-2, by administering one, or a small cocktail of, well-characterised, neutralising antibodies.

Patients recovering from COVID-19 have already been screened to identify neutralising antibodies, following analysis of relatively small numbers (100-500) of antibody sequences ^12,13^. A more extensive BCR repertoire analysis was performed on six patients in Stanford, USA with signs and symptoms of COVID-19 who also tested positive for SARS-CoV-2 RNA ^14^. Although no information was provided on the patient outcomes in that study, the analysis demonstrated preferential expression of a subset of immunoglobulin heavy chain (IGH) V gene segments with relatively little somatic hypermutation and showed evidence of convergent antibodies between patients.

To drive a deeper understanding of the nature of humoral immunity to SARS-CoV-2 infection and to identify potential therapeutic antibodies to SARS-CoV-2, we have evaluated the BCR heavy chain repertoire from 19 individuals at various stages of their immune response. We show that (1) there are stereotypic responses to SARS-CoV-2 infection, (2) infection stimulates both naïve and memory B cell responses, (3) sequence convergence can be used to identify putative SARS-CoV-2 specific antibodies, and (4) sequence convergence can be identified between different SARS-CoV-2 studies in different locations and using different sample types.

## Results

### COVID-19 disease samples

Blood samples were collected from n=19 patients admitted to hospital with acute COVID-19 pneumonia. The mean age of patients was 50.2 (SD 18.5) years and 13 (68%) were male. All patients had a clinical history consistent with COVID-19 and typical radiological changes.

Seventeen patients had a confirmatory positive PCR test for SARS-CoV-2. The patients experienced an average of 11 days (range 4-20) of symptoms prior to the day on which the blood sample was collected. Nine of the patients were still requiring hospital care but not oxygen therapy on day of sample collection (WHO Ordinal Scale Score 3), while eight were hospitalised requiring oxygen by conventional mask or nasal prongs (WHO Ordinal Scale Score 4) and two were hospitalised with severe COVID-19 pneumonia requiring high-flow nasal oxygen (WHO Ordinal Scale Score 5). On the day of sample collection, the direct clinical care team considered two patients to be deteriorating, four improving and the remaining thirteen were clinically stable.

### SARS-CoV-2 infection results in a stereotypic B cell response

IGHA and IGHG BCR sequencing yielded on average 135,437 unique sequences, and 23,742 clonotypes per sample (Supplementary Table 1). To characterise the B cell response in COVID-19, we compared this BCR repertoire data to BCR repertoire data from healthy controls obtained in a separate study ^15^. Comparing IGHV gene segment usage revealed a significantly different IGHV gene usage in COVID-19 patients compared to the healthy controls, most notably with increases in the usage of IGHV2-5 (2.6x IGHA, 1.0x IGHG increase), IGHV2-70 (4.6x IGHA, 4.1x IGHG increase), IGHV3-30 (2.0x IGHA, 1.4x IGHG increase), IGHV5-51 (3.5x IGHA, 2.0x IGHG increase), and IGHV4-34 (1.4x IGHA, 2.4x IGHG increase) in the COVID-19 patients (Figure 1A). All of these V gene segments have been previously observed in SARS-CoV-1 or SARS-CoV-2 specific antibodies^16^. IGHV4-34 has been shown to bind both autoantigens ^17^ and commensal bacteria ^18^ and has been associated with SLE ^19^. Our data extends this, showing that the proportion of sequences containing the autoreactive AVY & NHS sequence motifs within the IGHV region is significantly more frequent in improving COVID-19 patients compared to stable or deteriorating COVID-19 patients, specifically in the IGHG1 isotype subclass (p-value = 0.038; Supplementary Figure 2).

**Figure 1.**
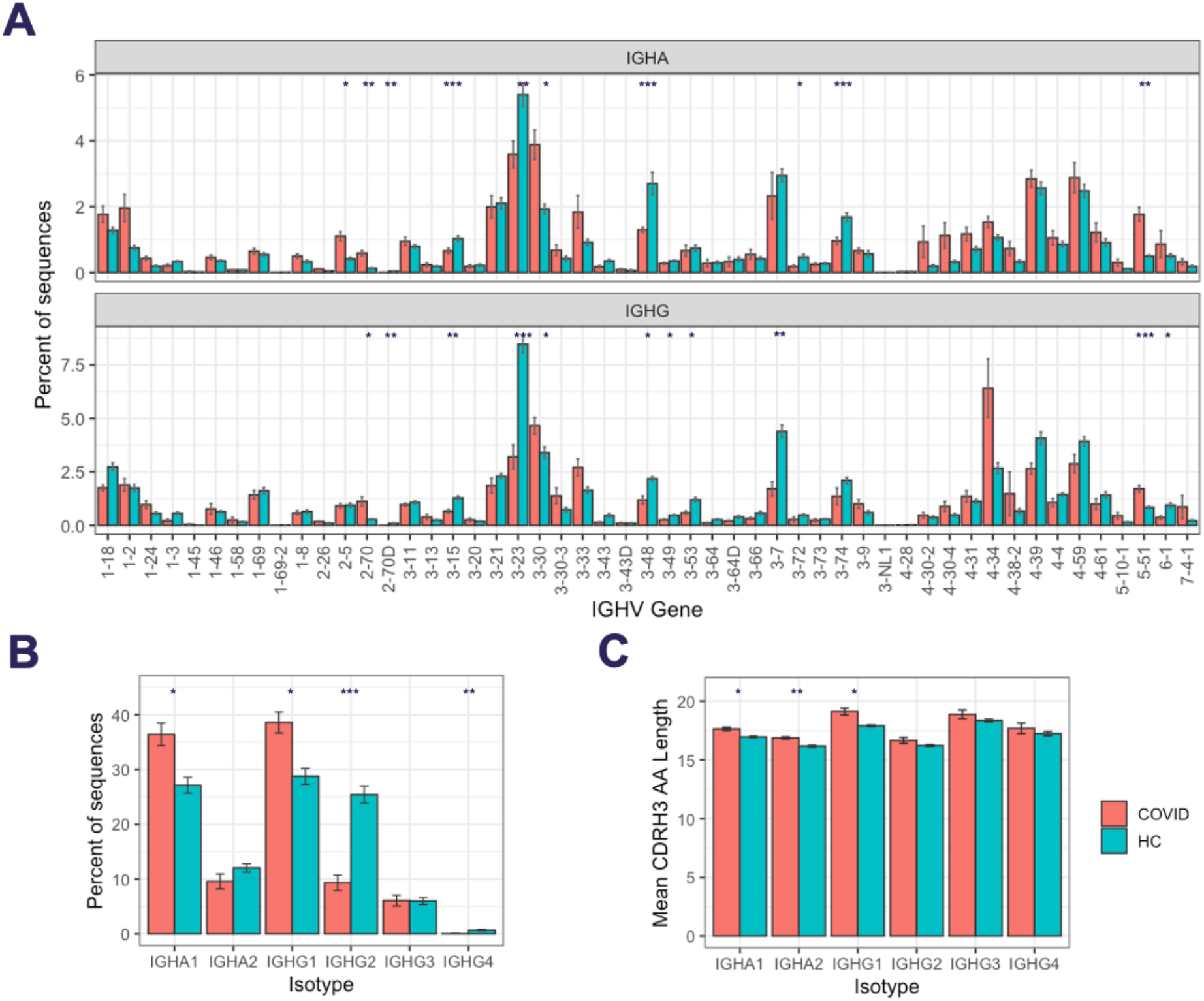
B cell responses to SARS-COV-2 infection. A) IGHV gene segment usage distribution per isotype subclass. B) Isotype subclass distribution between IGHA and IGHG subclasses, and C) mean BCR CDRH3 lengths from COVID-19 patients compared to healthy controls. For A-C, bars show mean values +/− standard error of the mean. Comparisons performed using t-tests, with adjusted p values using Bonferroni correction for multiple comparisons; * p < 0.05, ** p < 0.005, *** p < 0.0005.

Comparing isotype subclasses showed a significant increase in the relative usage of IGHA1 and IGHG1 in COVID-19 patients (Figure 1B) - these are the two first isotype subclasses that are switched to upon activation of IGHM ^20^. There was also an increase in the mean CDRH3 length of the BCRs in the COVID-19 patients, that was most pronounced in the IGHA1, IGHA2 and IGHG1 isotype subclasses (Figure 1C).

### SARS-CoV-2 infection stimulates both naïve and memory responses

To further investigate the COVID-19-specific B cell response, we analysed the characteristics of the BCR sequences that are consistent with recent B cell activation – somatic hypermutation, and clonal expansion. In healthy controls, for class-switched sequences, there is a clear unimodal distribution of sequences with different numbers of mutations, and a mean mutation count across IGHA and IGHG isotypes of 17.6 (Figure 2A). In the COVID-19 samples, the mean mutation count was 14.4, and there was a bimodal distribution with a separate peak of sequences with no mutations. This bimodal distribution was most pronounced in the IGHG1, IGHG3, and IGHA1 isotype subclasses, corresponding to the increased isotype usages. Such a distribution is consistent with an expansion of recently class-switched B cells that have yet to undergo somatic hypermutation. There was considerable variation between participants in the proportion of unmutated sequences (Supplementary Figure 1), which had no significant correlation with the number of days since symptom onset (R = 0.09, p = 0.72), but was lower in the deteriorating compared to improving patients (Figure 2B)

**Figure 2.**
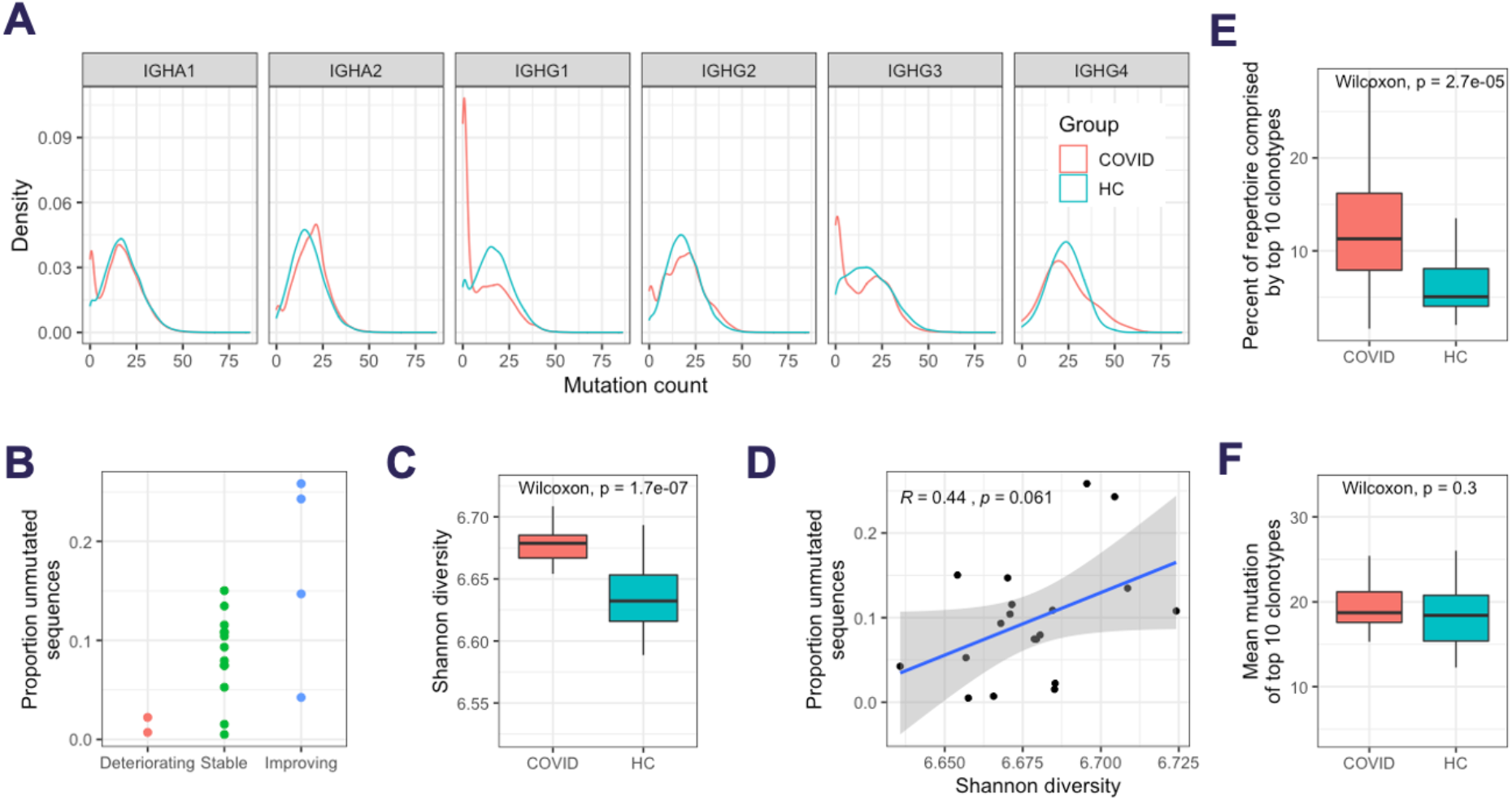
Response characteristics of SARS-CoV-2 infection. A) Distribution of sequences with different numbers of mutations from germline. B) Relationship between the proportion of the repertoire comprised by unmutated sequences, and the disease state C) Individual sequences were clustered together into related groups to identify clonal expansions (clonotypes). Diversity of all clonotypes in the repertoire calculated using the Shannon diversity index. To normalise for different sequence numbers for each sample, a random subsample of 1,000 sequences was taken. D) Correlation between the Shannon diversity index, and the proportion of unmutated sequences. E) The percent of all sequences that fall into the largest 10 clonotypes. F) Mean number of mutations of all sequences in the largest 10 clonotypes.

To investigate differential clonal expansion between patients, the Shannon diversity index of each repertoire was calculated (while accounting for differences in read depth through subsampling). A more diverse repertoire is indicative of a greater abundance of different clonal expansions. The BCR repertoires of the COVID-19 patients were significantly more diverse than the BCR repertoires of the healthy controls (Figure 2C); this increase in diversity was positively correlated with an increased proportion of unmutated sequences (R = 0.44, p = 0.061; Figure 2D). Interestingly, when we investigated the largest clonal expansions, despite having a more diverse repertoire, the largest clonal expansions in the COVID-19 samples were larger than in the healthy controls (Figure 2E). These large clonal expansions were also highly mutated and had similar levels of mutation between the COVID-19 samples and the healthy controls (Figure 2F).

### Sequence convergence can be used to identify putative SARS-CoV-2 specific antibodies

Given the skewing of the B cell response in the COVID-19 patients to specific IGHV genes, we next investigated whether the same similarity was also seen on the BCR sequence level between different participants. Such convergent BCR signatures have been observed in response to other infectious diseases ^21,^ and may be used to identify disease-specific antibody sequences.

Of the 435,420 total clonotypes across all the COVID-19 patients, 9,646 (2.2%) were shared between at least two of the participants (Figure 3A). As convergence could occur by chance or be due to an unrelated memory response from commonly encountered pathogens, the healthy control dataset was used to subtract irrelevant BCR sequences. Of the 9,646 convergent clonotypes, 1,442 (14.9%) were also present in at least one of the 40 healthy control samples. As expected, of the convergent clonotypes that were also present in the healthy control samples, the mean mutation count was significantly greater (Figure 3B), and the mean CDRH3 length significantly shorter (Figure 3C) than the convergent clonotypes that were unique to the COVID-19 patients.

**Figure 3.**
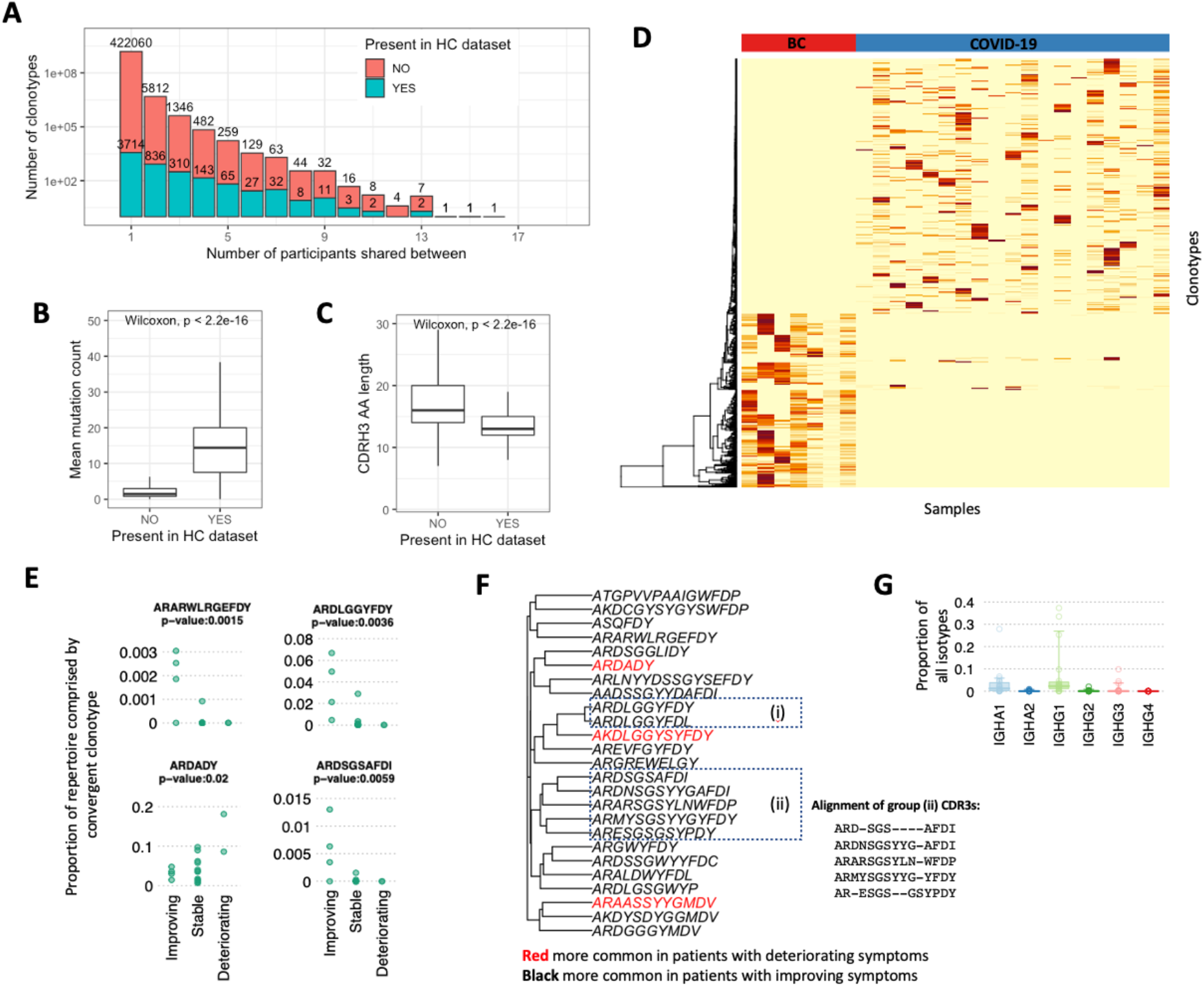
Convergent BCR sequence signature within individuals infected with SARS-CoV-2. A) Data from all patients and healthy controls were clustered together to identify convergent clonotypes. Shown is the number of clonotypes shared by different numbers of participants, grouped by whether the clonotypes are also present in the healthy control dataset. Of the convergent clonotypes, B) the mean mutation count, and C) the CDRH3 AA sequence length was compared between those that were convergent only within the SARS-CoV-2 patients, and those that were also convergent with the healthy control dataset. D) Heatmap of the 777 convergent COVID-19-associated clonotypes (observed between 4 or more COVID-19 participants) with the 469 convergent clonotypes from seven metastatic breast cancer (BC) patient biopsy samples, demonstrating that the convergent signatures are unique to each disease cohort. E) Percentage frequencies of four example convergent clonotypes grouped by clinical status. F) Similarity tree of convergent clonotype cluster centers that are significantly associated with clinical status. Groups (i) and (ii) indicate groups of similar convergent clonotypes. An alignment of group (ii) provided adjacent. G) Proportions of IGHA and IGHG of the convergent clonotypes that are associated with patients with improving symptoms.

To identify a set of SARS-CoV-2-specific antibody sequences with high confidence, we identified 777 convergent clonotypes that were shared between at least four of the COVID-19 patients, but not seen in the healthy controls. In parallel, for a comparison of convergent signatures, we performed the same analysis on a cohort of seven metastatic breast cancer patient biopsy samples ^22,^ which identified 469 convergent clonotypes. These convergent clonotypes were highly specific to each disease cohort (Figure 3D). The 777 COVID-19 convergent clonotypes had low mutation levels, with a mean mutation count of 2, and only 51 clonotypes with a mean mutation greater than 5. The sequences within the convergent clonotypes were primarily of the IGHG1 (70%) and IGHA1 (16%) subclasses (Supplementary Figure 3A). The convergent clonotypes used a diversity of IGHV gene segments, with IGHV3-30, IGHV3-30-3 and IGHV3-33 as the most highly represented (Supplementary Figure 3B). This IGHV gene usage distribution differs between that of the total repertoire, and IGHV3-30 is also the most highly used IGHV gene in the CoV-AbDab^16^.

We next tested whether these convergent clonotypes correlated with disease severity. Indeed, 25 of these convergent clonotypes were found to associate with clinical symptoms after correcting for multiple testing, of which 22 were observed at a significantly higher frequency in improving patients (Figure 3E and Supplementary Figure 4). This is a significantly higher proportion associated with clinical symptoms compared to that expected by chance (p-value = 0.018 by random permutations of labels). Interestingly, some of these clonotypes are common in patients comprising >0.1 % of a patient’s repertoire.

Furthermore, the convergent clonotypes that are associated with clinical symptoms cluster together (Figure 3F) and are found primarily in the IGHA1 and IGHG1 isotypes (Figure 3G).

### BCR clonotype sequence convergence signatures are shared between different COVID-19 studies in different locations and from different anatomical sites

To further explore whether the convergent clonotypes observed in our study were indeed disease specific, and to determine whether such convergence was common across studies and geographic regions, we compared these 777 convergent clonotypes to public B cell datasets.

First, we compared our data to RNAseq data of bronchoalveolar lavage fluid obtained from five of the first infected patients in Wuhan, China ^23^. These samples were obtained for the purpose of metagenomic analyses to identify the aetiological agent of the novel coronavirus but were re-analysed to determine whether we could extract any transcripts from BCRs. From the 10,038,758 total reads, we were able to identify 16 unique CDR3 AA sequences (

Supplementary Table 2). Of these, one had an exact AA match to a sequence in our data and shared the same V gene segment (IGHV3-15), and J gene segment (IGHJ4) usage (Figure 4A). The sequence had a CDRH3 AA length of 12, so such a match is unlikely to occur due to chance alone. The clonotype that the sequence belonged to contained 699 total sequences and was convergent between 8 of our 19 COVID-19 patients, but not present in the healthy controls. The clonotype was highly diverse, and the sequences had evidence of low mutation from germline, with a mean mutation count over all sequences of 4.8 (Supplementary Figure 5).

**Figure 4.**
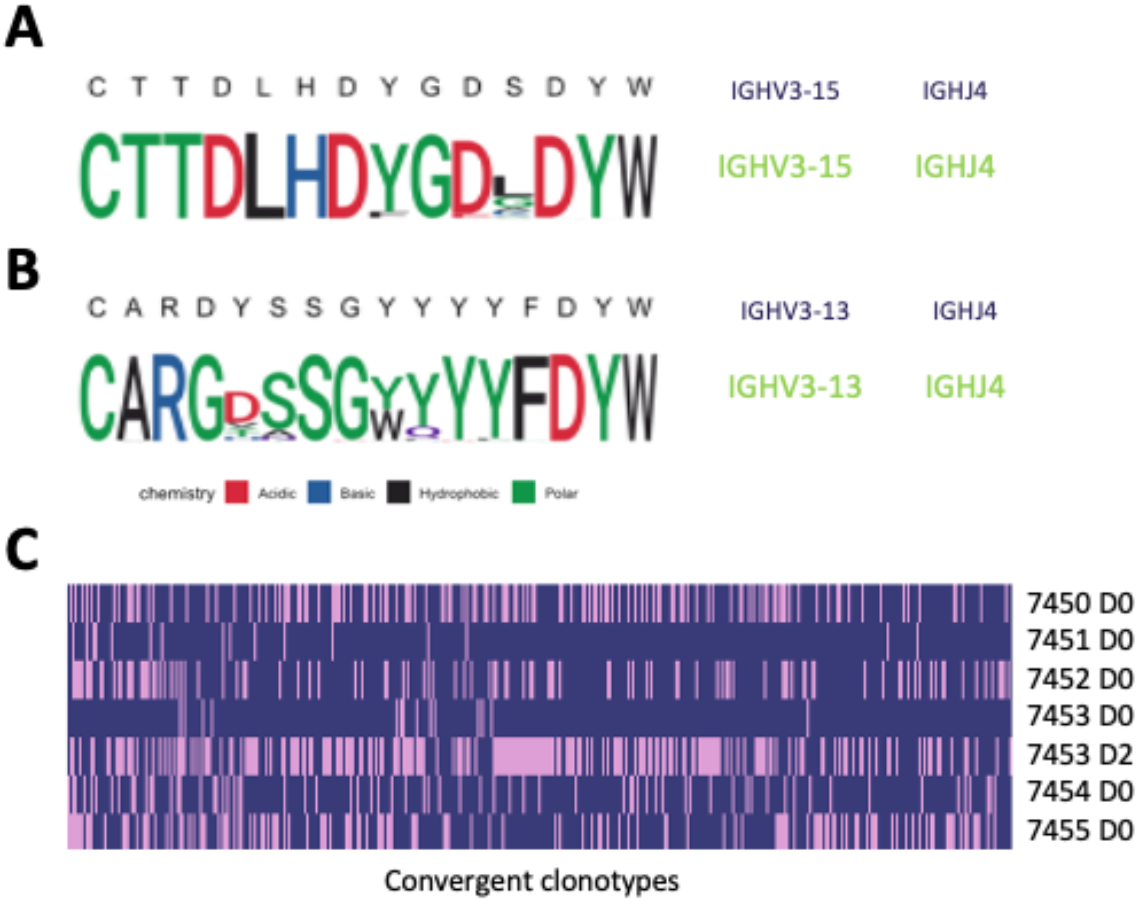
Matches of the 777 convergent clonotypes identified in the present study to other SARS-CoV-2 studies. A) CDRH3 sequence, and IGHV/IGHJ gene segments of a sequence identified in the bronchoalveolar lavage fluid of a SARS-CoV-2 patient from a Chinese cohort (shown across the top in black text), and a CDRH3 AA sequence logo unpacking the sequence diversity present in the convergent clonotype found in the COVID-19 patients in this study that had an exact AA match. B) CDRH3 sequence, and IGHV/IGHJ gene segment of an antibody in the CoV-AbDab (S304) that has SARS-CoV-1 and SARS-CoV-2 neutralising activity, alongside a CDRH3 AA sequence logo unpacking the sequence diversity in the convergent clonotype found in the COVID-19 patients in this study that had an exact AA match. C) Comparison of convergent clonotypes to the BCR data from Nielsen et al ^14^. Plotted along the x-axis are the 405 convergent clonotypes represented in at least one Nielsen et al. dataset. Each row represents a separate BCR repertoire from Nielsen et al.; pink shading indicates that the convergent clonotype has a match in the Nielsen dataset.

Next, we compared our 777 convergent clonotypes to CoV-AbDab – the Coronavirus Antibody Database [accessed 10^th^ May 2020] ^16^. At the time of access, this database contained 80 non-redundant CDRH3 sequences from published and patented antibodies proven to bind SARS-CoV-1 and/or SARS-CoV-2. We found 6 of our clonotypes to have high CDRH3 homology to the antibodies in CoV-AbDab (Figure 4B and Supplementary Figure 6). The most striking similarity was to S304, a previously described SARS-CoV-1 and SARS-CoV-2 receptor-binding domain antibody able to contribute to viral neutralisation ^24^. One of the 777 convergent clonotypes contained sequences with an exact CDRH3 AA sequence match and utilised the same IGHV and IGHJ germline gene segments to S304. This clonotype was convergent across 6 patients and had a mean mutation count of 1.1.

Finally, we compared our data to a publicly available BCR deep sequencing dataset from six COVID-19 patients from Stanford, USA. 405 of our 777 convergent clonotypes matched (using the same definition we used for clonotyping within our dataset) sequences in this dataset (Figure 4C), showing the high level of convergence between studies. The average number of clonotype matches to the Stanford COVID-19 patient repertoires was 95, but this varied considerably between patients and timepoints. Two of the six patients were seronegative at the day of sampling (7451 and 7453), and these two patients had the fewest clonotype matches (16 and 14 respectively). Patient 7453 had an additional sample taken two days later (following seroconversion), and at this point had a large increase in the number of clonotype matches to 204. There was one of the 777 convergent clonotypes that was found across all six of the Stanford patients, and 17 clonotypes that were found in at least four of five samples from seroconverted patients, but not found in the seronegative patients (Supplementary Table 3).

## Discussion

We have used deep sequencing of the BCR heavy chain repertoire to evaluate the B cell responses of 19 individuals with COVID-19. In agreement with previous studies, there was a skewing of the repertoire in the response to SARS-CoV-2 infection, with an increased use of certain V genes, an increase in the proportion of antibodies with longer CDRH3s, and an altered isotype subclass distribution ^14^. The significantly increased usage of IGHA1 observed in the COVID-19 patients is in line with mucosal responses, where the longer hinge in IGHA1 compared to IGHA2 may offer advantages in antigen recognition by allowing higher avidity bivalent interactions with distantly spaced antigens ^25^.

As anticipated, given the novel nature of the virus, that SARS-CoV-2 infection largely stimulated a characteristically naïve response, rather than a reactivation of pre-existing memory B cells – (1) there was an increased prevalence of unmutated antigen-experienced class-switched BCR sequences, (2) an increase in the diversity of class-switched IGHA and IGHG BCRs, and (3) an increase in the usage of isotype subclasses that are associated with recent viral immunity. These observations are consistent with an increase in the frequency of recently activated B cells in response to SARS-CoV-2. In addition to the naïve response, there was also evidence of a proportion of the response arising from memory recall. In the COVID-19 patients, the largest clonal expansions were highly mutated, equivalent to the level observed in healthy control cohort. Such a secondary response to SARS-CoV-2 has been previously observed ^26,^ and may be due to recall of B cells activated in response to previously circulating human coronaviruses, as recently highlighted^27,28^.

We observed a potential relationship between repertoire characteristics and disease state, with improving patients showing a tendency towards a higher proportion of unmutated sequences. The increased prevalence of autoreactive IGHV4-34 sequences in improving COVID-19 patients compared to stable or deteriorating COVID-19 patients potentially suggests a role for natural or autoreactive antibodies in resolving infection and lower risk of pathology. There is a clear need to expand on these findings by using larger sample cohorts and gathering more clinical data to aid understanding of the differences between individuals that respond with mild versus severe disease and have different recovery patterns. Building upon these observations could help to inform the future development of diagnostic assays to monitor and predict the progression of disease in infected patients.

A large number (777) of highly convergent clonotypes unique to COVID-19 were identified. Our approach of subtracting the convergent clonotypes also observed in healthy controls ^15,^ allowed us to identify convergence specific to the disease cohort. The unbiased nature of the BCR repertoire analysis approach means that, whilst these convergent clonotypes are likely to include many antibodies to the spike protein and other parts of the virus they may also include other protective antibodies, including those to host proteins.

Characterisation of the heavy chains we have identified, coupled with matched light chains to generate functional antibodies will permit analysis of the binding sites and neutralising potential of these antibodies. The report that plasma derived from recently recovered donors with high neutralising antibody titres can improve the outcome of patients with severe disease ^29,^ supports the hypotheses that intervention with a therapeutic antibody has the potential to be an effective treatment. A manufactured monoclonal antibody or combination of antibodies would also provide a simpler, scalable and safer approach than plasma therapy.

Sequence convergence between our 777 convergent clonotypes with heavy chains from published and patented SARS-CoV-1 and SARS-CoV-2 antibodies^16^ supports several observations. Firstly, it demonstrates that our approach of finding a convergent sequence signature is a useful method for enriching disease-specific antibodies, as we find matches to known SARS-CoV spike-binding antibodies. Secondly, it shows that the clonotypes observed in response to SARS-CoV-2 overlap with those to SARS-CoV-1, presumably explained by the relatively high homology of the two related viruses ^3^. Indeed, here we show that there is an overrepresentation of clonotypes that correlate with patient clinical symptoms than is expected by chance, and these BCR sequences are associated with the dominant IgA1 and IgG1 responses. Finally, it shows that the convergence extends beyond our UK COVID-19 disease cohort.

Further evidence for convergence extending beyond our disease cohort came from the comparisons of our 777 convergent clonotypes to deep sequencing datasets from China ^23^ and the USA ^14^. The dataset from the USA is also from BCR sequencing of the peripheral blood of COVID-19 patients, and here we found matches to 405 of our 777 clonotypes. The dataset from China was from total RNA sequencing of the bronchoalveolar lavage fluid of SARS-CoV-2 infected patients. Only 16 unique CDRH3 sequences could be identified in this whole dataset, but one of them matched a convergent clonotype in the current study, showing that convergence can be seen both between different locations, and different sample types. We believe that the identification of such high BCR sequence convergence between geographically distinct and independent datasets could be highly significant and validates the disease association of the clonotypes, as well as the overall approach.

In summary, our BCR repertoire analysis provides information on the specific nature of the B cell response to SARS-CoV-2 infection. The information generated has the potential to facilitate the treatment of COVID-19 by supporting diagnostic approaches to predict the progression of disease, informing vaccine development and enabling the development of therapeutic antibody treatments and prophylactics.

## Methods

### Clinical information gathering

Peripheral blood was obtained from patients admitted with acute COVID-19 pneumonia to medical wards at Barts Health NHS Trust, London, UK, after informed consent by the direct care team (NHS HRA RES Ethics 19/SC/0361). Venous blood was collected in EDTA Vacutainers (BD). Patient demographics and clinical information relevant to their admission were collected by members of the direct care team, including duration of symptoms prior to blood sample collection. Current severity was mapped to the WHO Ordinal Scale of Severity. Whether patients at time of sample collection were clinically Improving, Stable or Deteriorating was subjectively determined by the direct clinical team prior to any sample analysis. This determination was primarily made on the basis of whether requirement for supplemental oxygen was increasing, stable, or decreasing comparing current day to previous three days.

### Sample collection and initial processing

Blood samples were centrifuged at 150 x*g* for 15 minutes at room temperature to separate plasma. The cell pellet was resuspended with phosphate-buffered saline (PBS without calcium and magnesium, Sigma) to 20 ml, layered onto 15 ml Ficoll-Paque Plus (GE Healthcare) and then centrifuged at 400 x*g* for 30 minutes at room temperature without brake. Mononuclear cells (PBMCs) were extracted from the buffy coat and washed twice with PBS at 300 x*g* for 8 min. PBMCs were counted with Trypan blue (Sigma) and viability of >96% was observed. 5×10^6^ PBMCs were resuspended in RLT (Qiagen) and incubated at room temperature for 10 min prior to storage at –80°C. Consecutive donor samples with sufficient RLT samples progressed to RNA preparation and BCR preparation and are included in this manuscript.

Metastatic breast cancer biopsy samples were collected and RNA extracted as part of a previously reported cohort ^22^.

### RNA prep & BCR sequencing

Total RNA from 5×10^6^ PBMCs was isolated using RNeasy kits (Qiagen). First-strand cDNA was generated from total RNA using SuperScript RT IV (Invitrogen) and IgA and IgG isotype specific primers ^30^ including UMIs at 50 °C for 45 min (inactivation at 80 °C for 10 min).

The resulting cDNA was used as template for High Fidelity PCR amplification (KAPA, Roche) using a set of 6 FR1-specific forward primers ^30^ including sample-specific barcode sequences (6bp) and a reverse primer specific to the RT primer (initial denaturation at 95 °C for 3 min, 25 cycles at 98 °C for 20 sec, 60 °C for 30 sec, 72 °C for 1 min and final extension at 72 °C for 7 min). The amount of Ig amplicons (~450bp) was quantified by TapeStation (Beckman Coulter) and gel-purified.

Dual-indexed sequencing adapters (KAPA) were ligated onto 500ng amplicons per patient using the HyperPrep library construction kit (KAPA) and the adapter-ligated libraries were finally PCR-amplified for 3 cycles (98 °C for 15 sec, 60 °C for 30 sec, 72 °C for 30 sec, final extension at 72 °C for 1min). Pools of 10 and 9 libraries were sequenced on an Illumina MiSeq using 2×300 bp chemistry.

### Sequence processing

The Immcantation framework (docker container v3.0.0) was used for sequence processing ^31,32^. Briefly, paired-end reads were joined based on a minimum overlap of 20 nt, and a max error of 0.2, and reads with a mean phred score below 20 were removed. Primer regions, including UMIs and sample barcodes, were then identified within each read, and trimmed. Together, the sample barcode, UMI, and constant region primer were used to assign molecular groupings for each read. Within each grouping, usearch ^33,^ was used to subdivide the grouping, with a cutoff of 80% nucleotide identity, to account for randomly overlapping UMIs. Each of the resulting groupings is assumed to represent reads arising from a single RNA. Reads within each grouping were then aligned, and a consensus sequence determined.

For each processed sequence, IgBlast ^34^ was used to determine V, D and J gene segments, and locations of the CDRs and FWRs. Isotype was determined based on comparison to germline constant region sequences. Sequences annotated as unproductive by IgBlast were removed. The number of mutations within each sequence was determined using the shazam R package ^32^.

Sequences were clustered to identify those arising from clonally related B cells; a process termed clonotyping. Sequences from all samples were clustered together to also identify convergent clusters between samples. Clustering was performed using a previously described algorithm ^35^. Clustering required identical V and J gene segment usage, identical CDRH3 length, and allowed 1 AA mismatch for every 10 AAs within the CDRH3. Cluster centers were defined as the most common sequence within the cluster. Lineages were reconstructed from clusters using the alakazam R package ^36^. The similarity tree of the convergent clonontype CDR3 sequences was generated through a kmer similarity matrix between sequences in R.

### Public healthy control data processing

The healthy control BCR sequence dataset used here has been described previously ^15^. Only samples from participants aged 10 years or older, and from peripheral blood were used, resulting in a mean age of 28 (range: 11-51). Furthermore, only class-switched sequences were considered.

### Public SARS-CoV-2 bronchoalveolar lavage RNAseq data processing

The bronchoalveolar lavage data comes from a previously published study of SARS-CoV-2 infection ^23,^ with data available under the PRJNA605983 BioProject on NCBI. MIXCR v3.0.3 was used, with default settings, to extract reads mapping to antibody genes from the total RNASeq data ^37^.

### Public CoV-AbDab data processing

All public CDRH3 AA sequences associated with published or patented SARS-CoV-1 or SARS-CoV-2 binding antibodies were mined from CoV-AbDab^16,^ downloaded on 10^th^ May 2020. A total of 80 non-redundant CDRH3s were identified (100% identity threshold). These sequences were then clustered alongside the representative CDRH3 sequence from each of our 777 convergent clones using CD-HIT ^38,^ at an 80% sequence identity threshold (allowing at most a CDRH3 length mismatch of 1 AA). Cluster centres containing at least one CoV-AbDab CDRH3 and one convergent clone CDRH3 were further investigated.

### Public COVID-19 BCR sequence data processing

The fourteen MiSeq “read 1” FASTQ datasets from the six SARS-CoV-2 patients analysed in Nielsen et al.^14^ were downloaded from the Sequence Read Archive ^39^. IgBlast ^34^ was used to identify heavy chain V, D, and J gene rearrangements and antibody regions. Unproductive sequences, sequences with out-of-frame V and J genes, and sequences missing the CDRH3 region were removed from the downstream analysis. Sequences with 100% amino acid and isotype matches were collapsed. To circumvent the disparity in collapsed dataset sizes between pairs of replicates, we selected the replicate with the highest number of sequences for downstream analysis.

### Convergent Clonotyping Matching to Public Repertoires

The public SARS-CoV-2-positive^14^ and healthy control BCR repertoires^40^ were scanned for clonotype matches to our 777 convergent clonotype cluster centres. A BCR repertoire sequence was determined as a match if it had identical V and J genes, the same length CDRH3, and was within 1 AA mismatch per 10 CDRH3 AAs to a convergent clonotype representative sequence.

### Statistical analysis and graphing

Statistical analysis and plotting were performed using R ^41^. Plotting was performed using ggplot2 ^42^. Sequence logos were created using ggseqlogo ^43^. Specific statistical tests used are detailed in the figure legends. Correlations of IGHV4-34 autoreactive motifs and convergent clonotypes was performed by manova in R.

## Acknowledgments

The authors would like to first and foremost acknowledge and thank the patients who consented to providing their samples to this study. We are grateful to Ursula Arndt, and the RTD team at Illumina who performed the sequencing reactions to generate data for analysis. We also thank Felicia Anna Tucci (University of Oxford) for her help with the BCR sequencing methods.

## Author contributions

All authors discussed methodology, results and contributed to the final manuscript. JO, OC, AL, GT, DP, PP conceived and designed the study. JDG, SS, MIJR, RJMB-R and AK conducted the data analysis with input from JO, RM, JD, GJK, DKF, LJ, RJMB-R, CMD, OC, W-YJL, GT, PP and DP. GT, PP, DP recruited COVID-19 participants and executed the clinical protocols. SS, JD and W-YJL processed the COVID-19 clinical samples. CC and JS recruited the breast cancer participants. SS and JD processed the breast cancer samples. JB oversaw sequencing of the libraries. JDG, JO, GJK, SS and RM wrote the manuscript with input from all co-authors. All authors read and approved the final manuscript.

## Data availability

COVID-19 BCR sequence data will be made available upon publication.

## Supplementary information

**Supplementary Figure 1.**
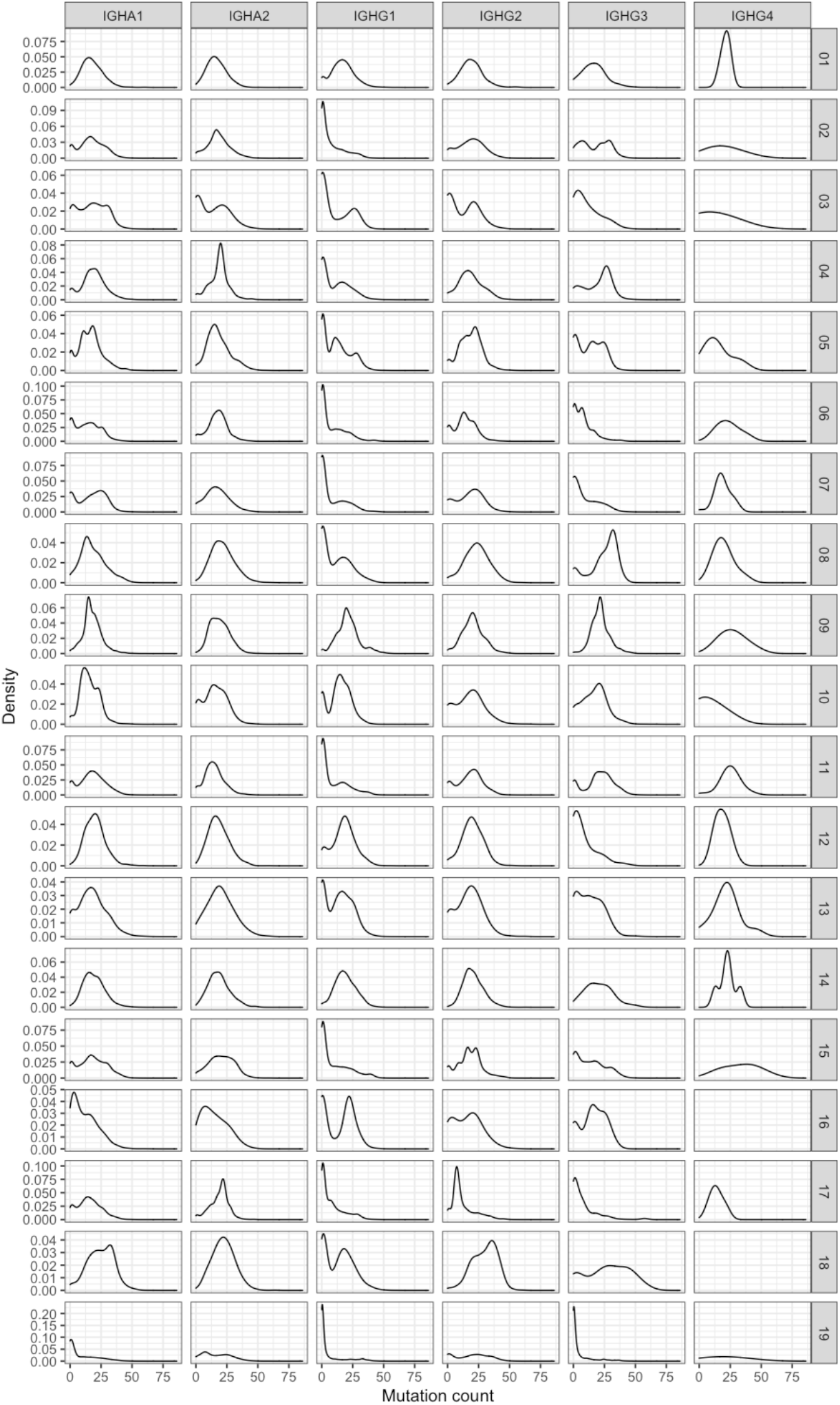
Distribution of sequences with different numbers of mutations from germline. Each row is a different COVID-19 patient and (right)

**Supplementary Figure 2.**
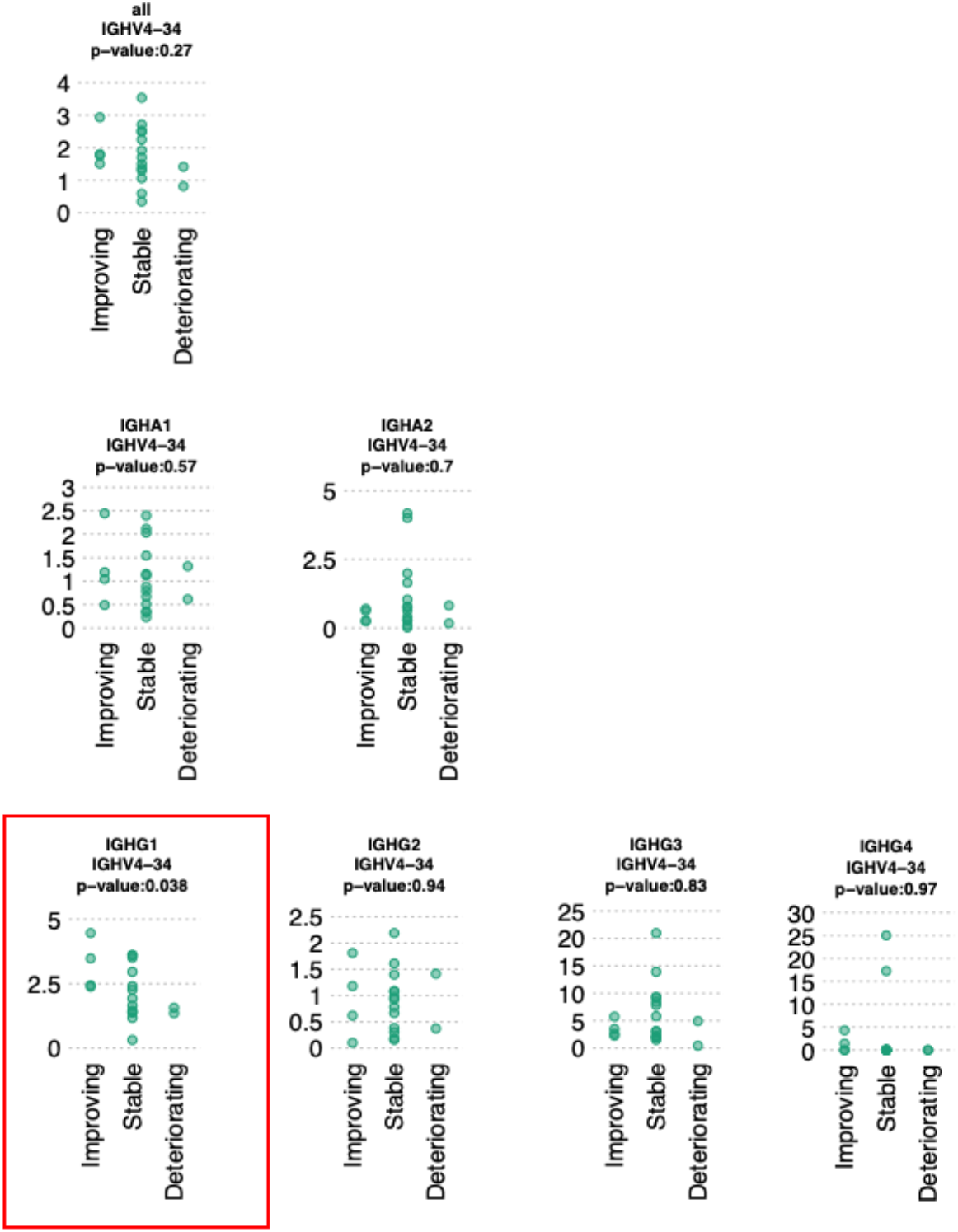
The proportion of IGHG1 sequences containing the autoreactive “NHS and “AVY” motifs between COVID patients with improving, stable or worsening symptoms. IGHG1 (red box) was the only significant correlation. P-values are determined by ANOVA.

**Supplementary Figure 3.**
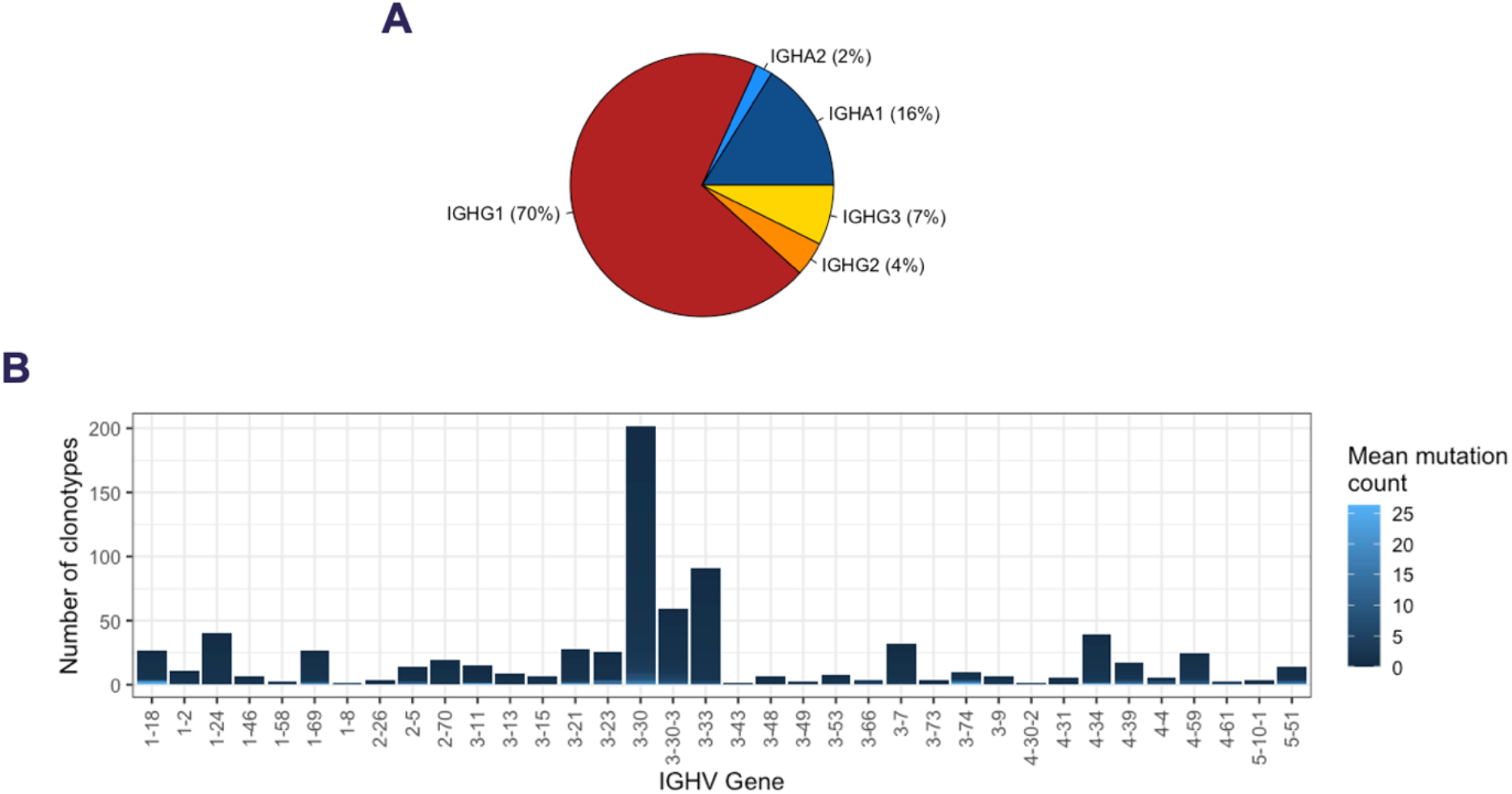
Properties of the 777 convergent clonotypes A) Isotype subclass usage of the sequences with the 777 convergent clonotypes. B) IGHV gene segment usage of the 777 convergent clonotypes.

**Supplementary Figure 4.**
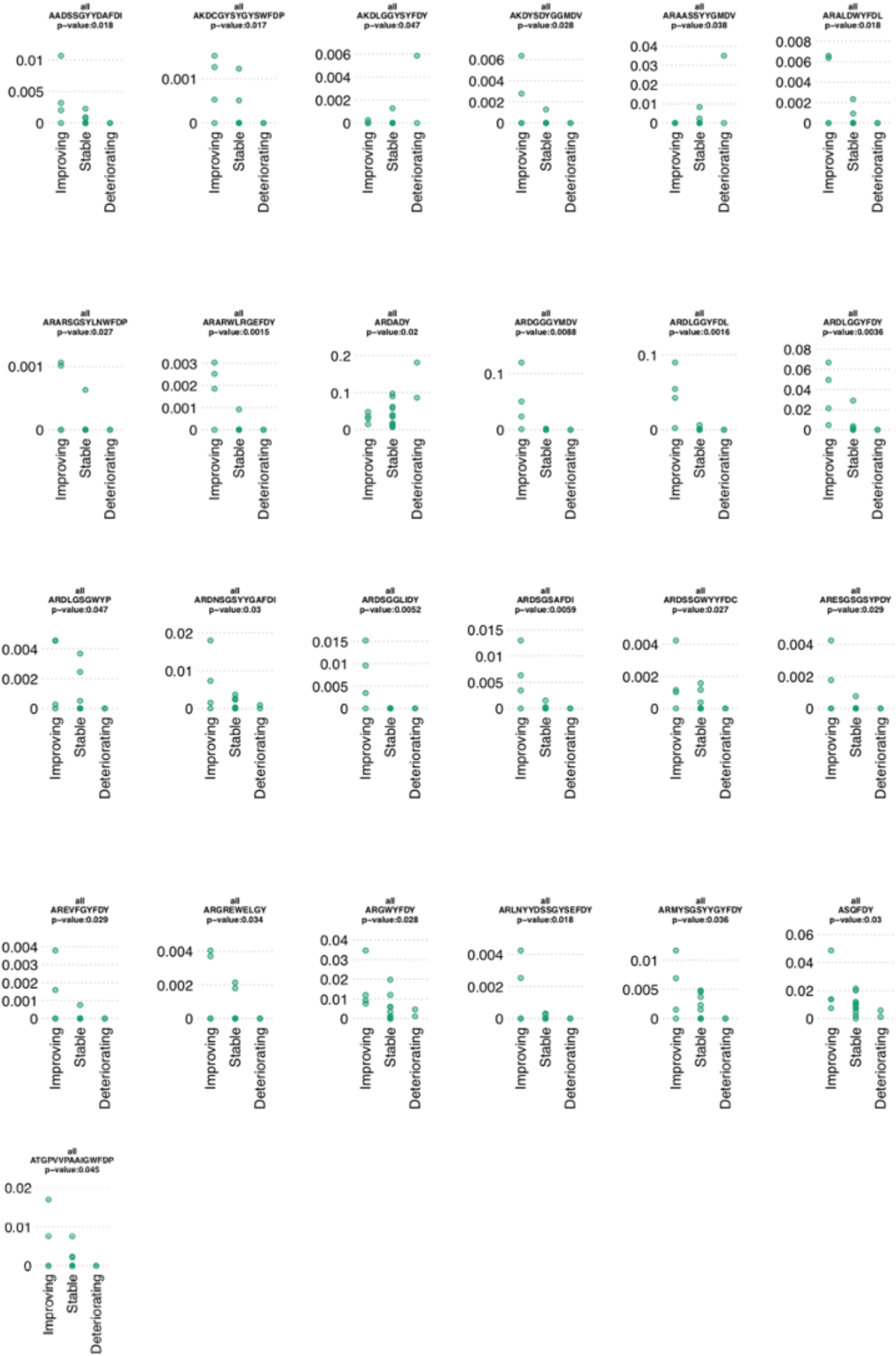
Percentage frequencies of the convergent clonotypes grouped by clinical status that significantly associated with clinical status.

**Supplementary Figure 5.**
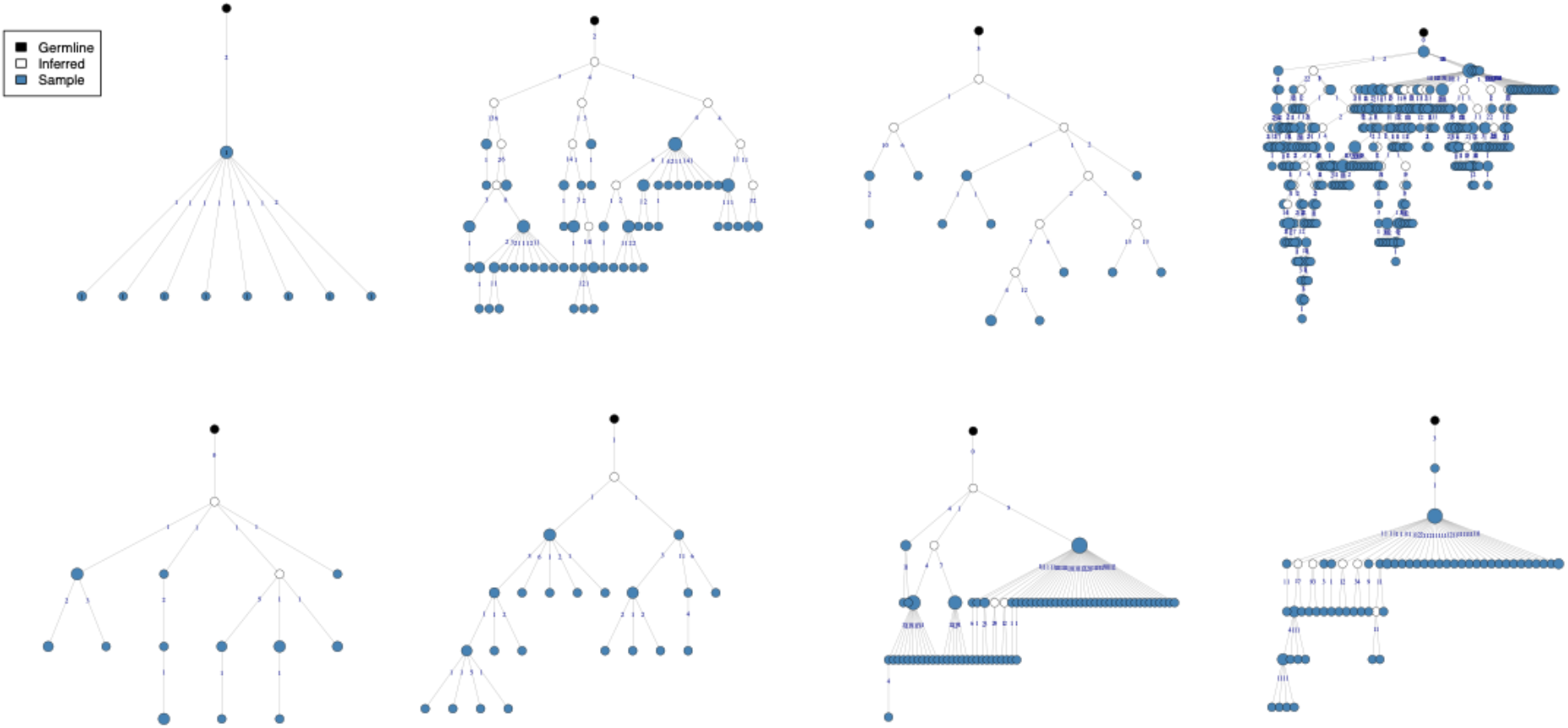
Lineage trees of the convergent clonotype that matched to the bronchoalveolar lavage fluid data. Each lineage tree represents the members of the clonotype from each of the eight patients it was present in. Each node represents a unique sequence within the clonotype lineage tree, with the size indicative of the number of duplicate sequences present. Numbers on the edges of adjoining nodes show the number of mutations between the sequences.

**Supplementary Figure 6.**
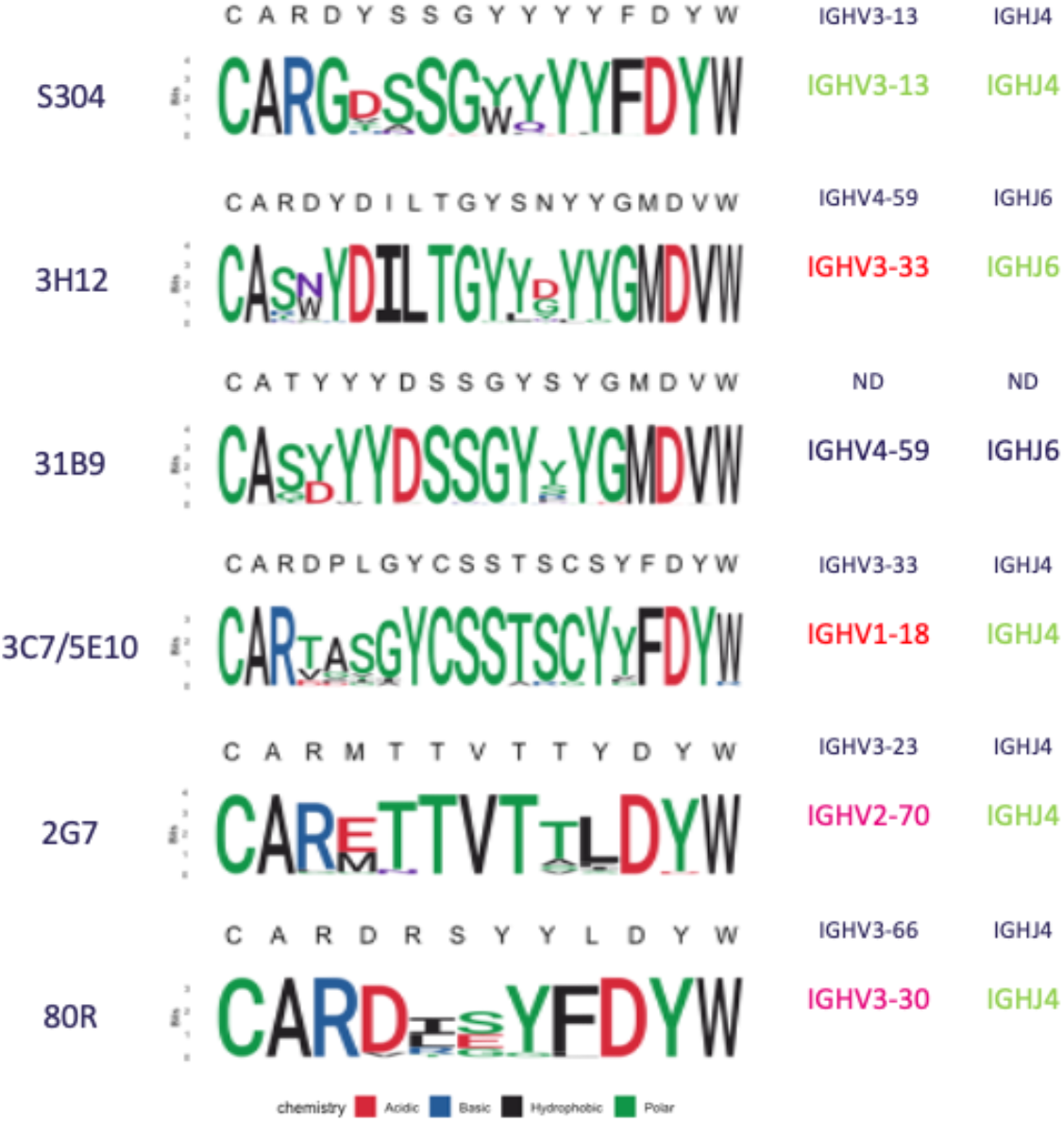
Logo plots unpacking the sequence diversity present for the convergent clonotypes that clustered with CoV-AbDab SARS-CoV-1 or SARS-CoV-2 binding antibodies. The CoV-AbDab reference CDRH3 and IGHV/IGHJ gene segment is displayed above each Logo plot. Gene transcript matches are shown in green, while mismatches are shown in red. The full sequence for 31B9 is not yet publicly available, so its genetic origins are not determined (ND).

**Supplementary Table 1.**
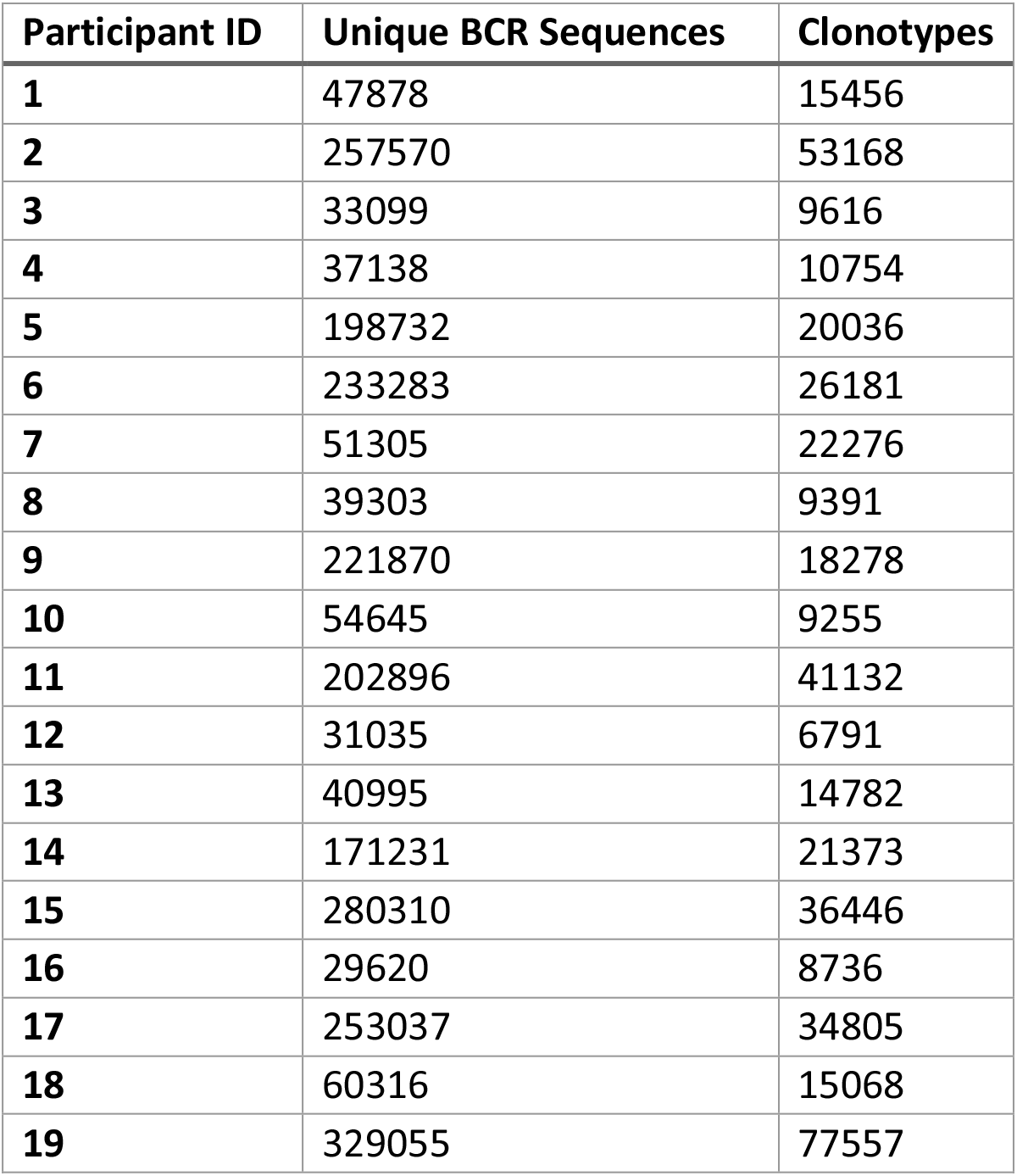
Summary of number of unique sequences, and number of clonotypes obtained for each COVID-19 patient.

| Participant ID | Unique BCR Sequences | Clonotypes |
| --- | --- | --- |
| 1 | 47878 | 15456 |
| 2 | 257570 | 53168 |
| 3 | 33099 | 9616 |
| 4 | 37138 | 10754 |
| 5 | 198732 | 20036 |
| 6 | 233283 | 26181 |
| 7 | 51305 | 22276 |
| 8 | 39303 | 9391 |
| 9 | 221870 | 18278 |
| 10 | 54645 | 9255 |
| 11 | 202896 | 41132 |
| 12 | 31035 | 6791 |
| 13 | 40995 | 14782 |
| 14 | 171231 | 21373 |
| 15 | 280310 | 36446 |
| 16 | 29620 | 8736 |
| 17 | 253037 | 34805 |
| 18 | 60316 | 15068 |
| 19 | 329055 | 77557 |

**Supplementary Table 2.**
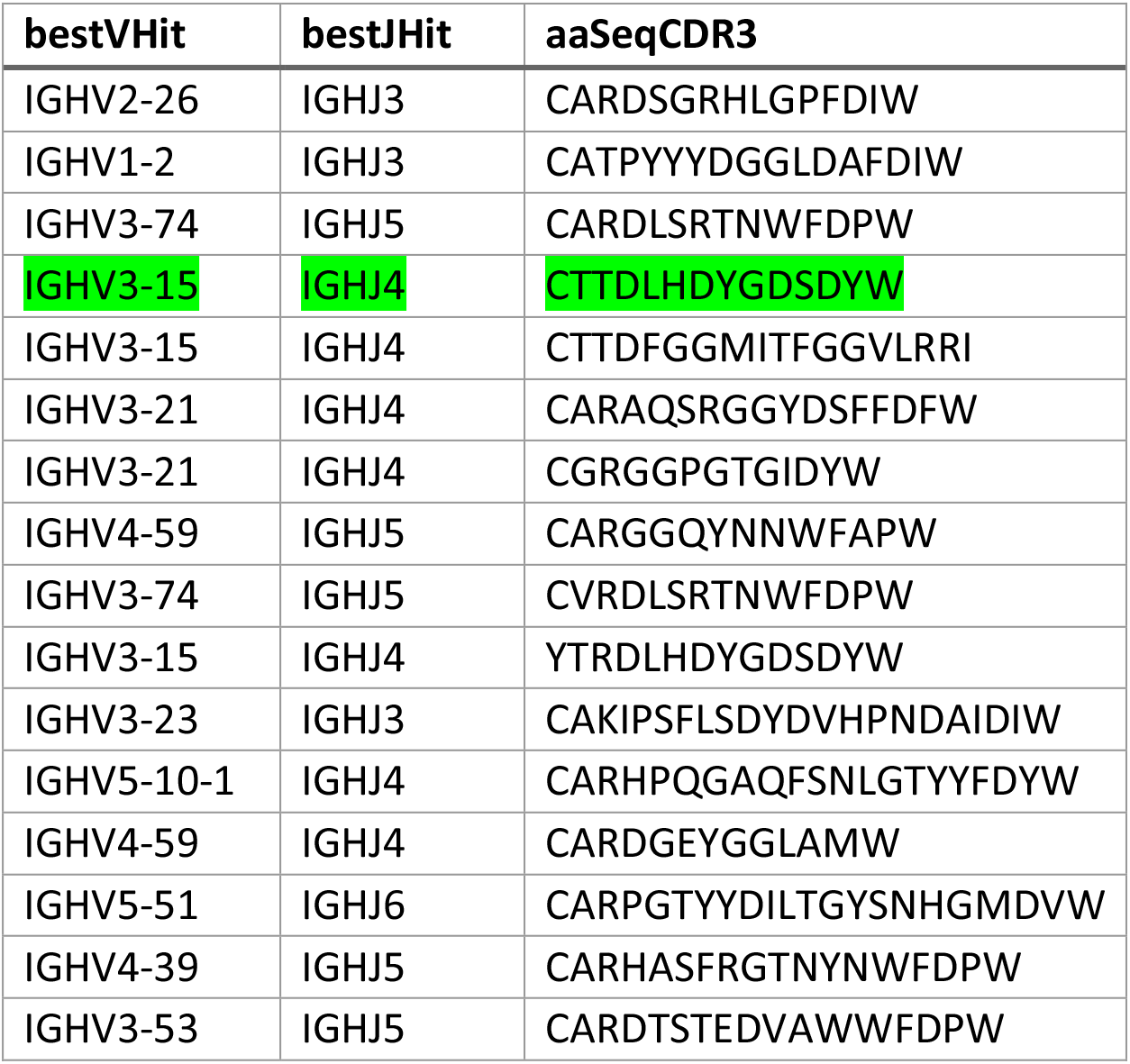
CDRH3 AA sequences identified from bronchoalveolar RNAseq data. Highlighted in green is the one identified in our SARS-CoV-2 patient dataset.

**Supplementary Table 3.** A subset of the 777 convergent clonotypes that matched to at least 4 of the samples in the Nielsen et al ^14^ data.

| Cluster Center CDRH3 | IGHV gene | IGHJ gene | Total Matches | Seropositive Matches | Seronegative Matches |
| --- | --- | --- | --- | --- | --- |
| ARGFDP | IGHV4-34 | IGHJ5 | 5 | 5 | 0 |
| ARVFDY | IGHV3-30 | IGHJ4 | 5 | 4 | 1 |
| ARELSYYGMDV | IGHV3-30 | IGHJ6 | 5 | 4 | 1 |
| AREDYGDYGFDY | IGHV3-30 | IGHJ4 | 5 | 4 | 1 |
| ARGFDH | IGHV4-61 | IGHJ4 | 4 | 4 | 0 |
| ARDSGGLIDY | IGHV3-30 | IGHJ4 | 4 | 4 | 0 |
| AREVMVYLDY | IGHV3-33 | IGHJ4 | 4 | 4 | 0 |
| ARDSGSAFDI | IGHV3-30-3 | IGHJ3 | 4 | 4 | 0 |
| ANDLYYGMDV | IGHV3-30 | IGHJ6 | 4 | 4 | 0 |
| AREGPDAFDI | IGHV1-18 | IGHJ3 | 4 | 4 | 0 |
| AKEGIVAFDY | IGHV3-30 | IGHJ4 | 4 | 4 | 0 |
| ARQEHYYYGMDV | IGHV5-51 | IGHJ6 | 4 | 4 | 0 |
| ARPYSGSYRGYFDY | IGHV3-30 | IGHJ4 | 4 | 4 | 0 |
| ARSRGGSYYGFDY | IGHV3-30 | IGHJ4 | 4 | 4 | 0 |
| ARDLDYYDSSGFDY | IGHV3-7 | IGHJ4 | 4 | 4 | 0 |
| AKARGGSYLDAFDI | IGHV3-30 | IGHJ3 | 4 | 4 | 0 |
| ARVDYYDSSGYYRDY | IGHV1-69 | IGHJ4 | 4 | 4 | 0 |
| TTGTWYYDSSGYSNDAFDI | IGHV3-15 | IGHJ3 | 4 | 4 | 0 |
| ARGIDY | IGHV3-23 | IGHJ4 | 4 | 3 | 1 |
| ARDLGDYGMDV | IGHV3-53 | IGHJ6 | 4 | 3 | 1 |

